# Resting-state and task-evoked phase-amplitude coupling between cardiac and neural rhythms is sensitive to anxiety severity

**DOI:** 10.64898/2026.08.06.743330

**Authors:** Asa Young, Jonathan W. Schooler

## Abstract

**Background:** High-frequency heart-rate variability (HF-HRV) is a downstream marker of cortico-autonomic regulation linked to executive function. A recent proposal suggests it indexes regulatory processes because it is both a product of and contributor to neural organization within the prefrontal cortex. The oscillatory phase of HF-HRV has been reported to modulate fronto-central EEG amplitude at rest, and this coupling is attenuated in individuals with schizophrenia relative to healthy controls. We evaluated this brain–body correspondence in relation to anxiety symptomatology, which is likewise associated with autonomic dysregulation.

**Methods:** Concurrent EEG and electrocardiography (ECG) were recorded in 22 nonclinical adults at rest and during a mental arithmetic task. Participants rated anxiety severity using the Generalized Anxiety Disorder 7-item scale (GAD-7). We evaluated between-person associations between anxiety severity and HF-HRV–EEG coupling, as well as within-person differences in coupling between rest and mental arithmetic.

**Results:** Anxiety symptomatology was associated with decreased resting-state phase-amplitude coupling between HF-HRV phase and fronto-central theta amplitude, independent of EEG and HRV covariates. Relative to rest, HF-HRV–theta coupling increased during a cognitive task, independent of condition-related changes in EEG, HR, or respiration. Anxiety severity moderated the task-evoked changes in heart-brain coupling such that more anxious individuals exhibited larger condition-related differences. Simple-slope analyses indicated anxiety’s effect on coupling at rest was absent when engaged in a task.

**Conclusions:** HF-HRV–theta phase-amplitude coupling captured anxiety-related and state-dependent variance not evident in conventional cardiac or neural indices. This coupling may index a state-sensitive component of cortico-autonomic regulation, although its putative functional role requires direct testing.

---

Heart rate variability (HRV), fluctuations in the interval between consecutive heartbeats, reflects the sympathetic and parasympathetic influences on cardiac activity. High-frequency HRV (HF-HRV; 0.15-0.40 Hz) is commonly interpreted as an index of vagally mediated parasympathetic regulation closely associated with respiratory sinus arrhythmia—the recurrent cardiac acceleration at inspiration and deceleration at expiration (Shaffer & Ginsberg, 2017). Greater HF-HRV reflects flexible autonomic regulation and has accordingly been linked to a range of clinically relevant outcomes, including reduced worry and rumination (Ottaviani et al., 2016), lower cardiovascular risk (Chalmers et al., 2014), and regulated emotional responding (Appelhans & Luecken, 2006). Although HF-HRV is commonly treated as a downstream marker of cortico-autonomic regulation, heart-rate oscillations have been proposed to contribute to the organization of neural activity within prefrontal systems involved in emotion regulation (Mather & Thayer, 2018). In this view, HRV may index regulatory processes because it is itself a product of and contributor to cognitive and emotional regulation.

A growing body of work supports the broader view that bodily rhythms modulate neural excitability and stimulus processing in functionally meaningful ways, contributing not only to homeostasis but also to perception, emotion, and cognition (Azzalini et al., 2019; Young et al., 2022; Young et al., 2026). Consistent with this view, Raut et al. (2021) demonstrated that the large-scale organization of resting-state functional connectivity can be parsimoniously accounted for by global traveling waves of excitability linked to peripheral indices of arousal. Peripheral physiological rhythms may therefore both reflect and contribute to variation in global brain states.

One mechanism by which such peripheral rhythms may shape cortical dynamics is phase-amplitude coupling, a form of cross-frequency coupling in which the oscillatory phase of a low-frequency rhythm modulates (nests) the amplitude of a comparatively higher-frequency rhythm (Tort et al., 2008). Consistent with the proposal that HF-HRV may contribute to the organization of prefrontal neural activity (Mather & Thayer, 2018), Sargent et al. (2024) reported phase-amplitude coupling (PAC) between the phase of HF-HRV and the amplitude of all five characteristic electroencephalographic (EEG) frequency bands in fronto-central channels^1^. Granger causality indicated a predominantly ascending relationship, with stronger heart-to-brain than brain-to-heart effects. HF-HRV–EEG phase-amplitude coupling may therefore provide a candidate mechanism linking heart rate variability to clinically relevant outcomes.

Clinical disorders involving cognitive and affective dysregulation, including post-traumatic stress disorder (Ge et al., 2020), major depression (Koch et al., 2019), and schizophrenia (Haigh et al., 2021), are often associated with reduced HF-HRV, suggesting that atypical autonomic regulation may accompany, and perhaps contribute to, higher-order dysfunction. If HRV–EEG phase-amplitude coupling indexes regulatory processes, it may likewise be altered in conditions characterized by autonomic and cognitive dysregulation. Consistent with this possibility, Sargent et al. (2025) reported reduced frontal theta and alpha HRV–EEG phase-amplitude coupling in individuals with first-episode schizophrenia relative to healthy controls. Coupling uniquely predicted group membership, whereas HF-HRV and EEG power alone did not.

Heart-brain coupling varies according to traits and states, but the manner of its variation may provide insights regarding its function. For example, the heartbeat-evoked potential (HEP), an event-related potential time-locked to the R-wave, is an index of cortical processing of cardiac afference (Schandry et al., 1986). Consistent with this interoceptive interpretation, HEP amplitude is associated with interindividual differences in heartbeat counting accuracy (Pollatos & Schandry, 2004) and increases during interoceptive attention (Petzschner et al., 2019; Coll et al., 2021). It is enhanced, relative to healthy controls, in clinical conditions involving internal hypervigilance, including obsessive-compulsive disorder (Yoris et al., 2017) and generalized anxiety disorder (Pang et al., 2019). By contrast, information-theoretic measures of statistical dependence^2^ between cardiac and low-frequency EEG phase decrease with task disengagement and declining vigilance during a non-interoceptive sustained-attention task—an effect not mirrored by HEP amplitude (Corcoran et al., 2025). Thus, distinct patterns of trait- and state-related variation may help differentiate heart–brain coupling associated with interoceptive processing from coupling associated with non-interoceptive faculties.

Anxiety is a compelling target for evaluating HRV–EEG phase-amplitude coupling because it is characterized by autonomic and attentional dysregulation, including persistent arousal, hypervigilance, and subjective distress (Thayer & Lane, 2000). It also provides a useful contrast for interpreting the functional significance of HRV–EEG phase-amplitude coupling. Schizophrenia has been associated with interoceptive deficits (Koreki et al., 2021; Fan et al., 2025), whereas anxiety has been linked to enhanced interoceptive processing (Young et al., 2026). If HRV–EEG phase-amplitude coupling primarily indexes interoceptive processing, anxiety symptomatology may therefore be associated with increased coupling. Conversely, if coupling reflects coordination between cardiac and neural rhythms involved in a non-interoceptive facet of cognitive regulation, anxiety symptomatology may be associated with decreased coupling. Given prior evidence that HRV–EEG coupling is reduced in theta and alpha bands in first-episode schizophrenia (Sargent et al., 2025), we asked whether coupling in these same bands tracks variation in self-reported anxiety symptoms within a nonclinical sample. We examined this relationship at rest and during mental arithmetic to distinguish anxiety-related variation in spontaneous coupling from variation in task-evoked coupling.

## Methods and Materials

### Participants

24 nonclinical individuals were recruited from the student population of the University of California, Santa Barbara. Eligibility criteria excluded individuals with a history of cardiorespiratory, gastrointestinal, or neuropsychiatric disorders, as well as traumatic brain injury. 2 participants were excluded for exhibiting a resting-state root mean square of successive differences (RMSSD), a time-domain index of heart-rate variability, in excess of 2.5 standard deviations^3^, yielding a final sample of 22 participants (10 males) with a mean age of 24.64 ± 4.48 years. The sample completed the Generalized Anxiety Disorder 7-item (GAD-7; Spitzer et al., 2006), a brief self-report questionnaire assessing general anxiety severity over the past 2 weeks. The sample reported a mean GAD-7 score of 5.82 ± 4.25, indicating anxiety symptoms ranging from minimal to moderate.

### Data Collection

These data were drawn from a larger multimodal psychophysiological study comprising resting-state, mental arithmetic, and meditation conditions administered in counterbalanced order. The present analyses included electroencephalography (EEG), electrocardiography (ECG), and respiratory effort data from the resting-state and mental arithmetic conditions.

EEG, ECG, and respiratory effort were recorded simultaneously while participants completed, in counterbalanced order, a 15-minute resting-state condition and a 5-minute mental arithmetic task. During rest, participants maintained a relaxed posture while staring at a fixation cross and engaging in undirected thought. During arithmetic, participants maintained the same posture while serially subtracting 7s from 1000. ECG was acquired at 1,000 Hz with a BIOPAC MP160 system equipped with ECG100C (see footnote^4^ for module settings) and RSP100C modules (see footnote^5^ for module settings). ECG was captured using two BIOPAC EL503 electrodes placed on either side of the left chest cavity and one EL503 ground electrode placed on the right rib cage. Respiratory effort was recorded using a BIOPAC TSD201 respiratory effort transducer—an elastic velcro band tightly strapped around the participants’ abdomen. EEG was recorded using a standard 19-channel Electro-Cap International cap (10-20 system) sampled at 1,000 Hz with a TMSi Mobita wireless amplifier. BIOPAC Systems AcqKnowledge 5.0 data acquisition and analysis software was employed for acquiring bodily and EEG signals.

To achieve temporal alignment between ECG and EEG series, both amplifiers monitored a light-sensitive diode affixed to the participant’s screen. One-second black-and-white flashes, hidden from the participant, were presented to the diode at the beginning and end of each condition and every 10 seconds, thereby anchoring the separate series to a common temporal reference frame. These pulses were used to align and epoch the bodily and EEG signals offline using in-house MATLAB scripts.

### Pre-Processing

Data were exported from AcqKnowledge 5.0 as MATLAB (2023a) files. EEG recordings were imported into EEGLAB (Delorme & Makeig, 2004) for preprocessing. Raw data were bandpass filtered 0.5-50 Hz, excluding the light-sensitive channel. Signals were re-referenced to the average, after which channel data and power spectra were inspected to identify dead or noisy electrodes. Bad channels were removed prior to independent component analysis (ICA), which used the default extended Infomax settings with sphering enabled. ICA components were automatically classified using ICLabel (Pion-Tonachini et al., 2019) and the time series, power spectra, and spatial topography of each component were visually inspected to reject noise components. Component rejection targeted eye, muscle, and cardiac artifacts. Following component rejection, rejected channels were interpolated using a spherical spline interpolation over the scalp.

### HF-HRV Oscillatory Phase Estimation

Following initial EEG preprocessing, HF-HRV phase time series were generated from the ECG signal. R-waves were identified using the Pan-Tompkins algorithm (Pan & Tompkins, 1985; Fig. 1A) implemented in MATLAB by Sedghamiz (2014). To ensure detection quality, a two-stage guardrail procedure was applied. First, competing detections (e.g., T-wave detection) within a 300 ms window were resolved by retaining only the peak of highest amplitude. Second, the resulting inter-beat intervals (IBIs) were screened against physiological bounds of 400-1500 ms; within-window duplicates that survived the first stage were again arbitrated by amplitude, while implausibly long intervals were retained and flagged for visual inspection. The indices of accepted R-waves were plotted against the raw ECG for visual inspection. Accepted R-waves were used to construct a sample-level IBI series (in ms) by assigning each interval value to all samples between consecutive R-peaks. Samples preceding the first detected R-peak and following the last were excluded and corresponding EEG time points were likewise excluded from downstream analysis.

**Figure 1.**
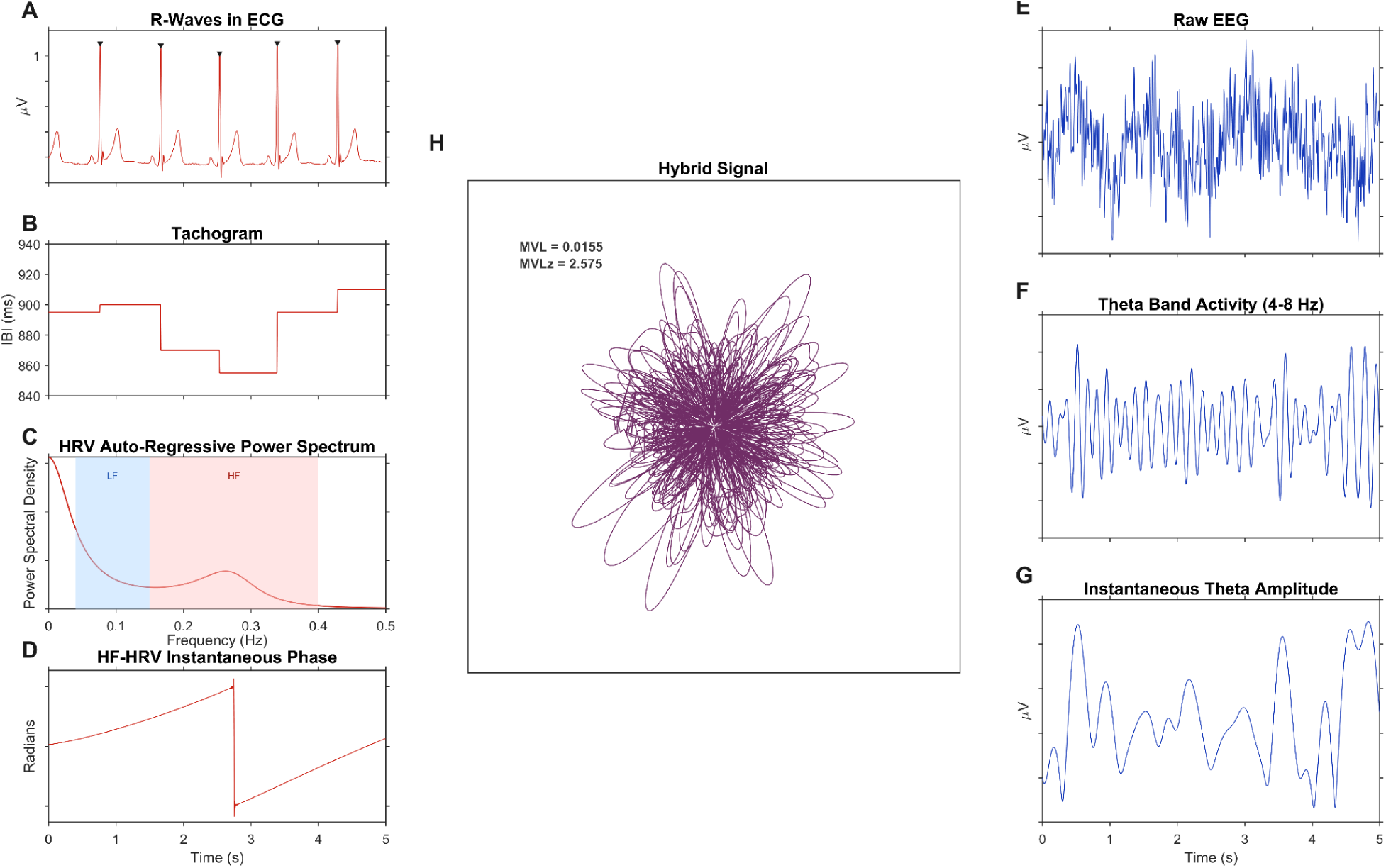
Signal Processing Methods in a Representative Participant. *Note. Figures A-D illustrate the methods for estimating participant-specific HF-HRV instantaneous phase angles. Figures E-G illustrate the methods for estimating instantaneous EEG band (e.g. theta) amplitude. Figure H depicts the hybrid signal used to quantify phase-amplitude coupling. HF-HRV phase provides the angular coordinate, and the instantaneous amplitude of the higher-frequency signal provides the radial coordinate, such that the resulting polar representation is plotted in Cartesian space. HF-HRV instantaneous phase is derived by first identifying R-spike indices using the Pan-Tompkins algorithm (A), creating a sample-level tachogram of interbeat intervals (B), and surveying the power spectral distribution of HRV through autoregressive means (C). The sample-level tachogram is then filtered within ± 0.05 Hz of the HF-HRV peak, and instantaneous phase is extracted by taking the angle of the Hilbert transform (D). Instantaneous amplitude of, for example, the theta band is derived by filtering artifact-free EEG (E) to the characteristic theta frequency band (F). Instantaneous amplitude is generated by taking the absolute value of the Hilbert transform of this filtered signal (G)*.

The IBI series was resampled to 4 Hz using cubic spline interpolation, linearly detrended, and submitted to autoregressive (AR) spectral estimation via the Yule-Walker equations. The AR model order was selected from the candidate range of 8-20 in accordance with Malik (1996) recommendations, with the optimal order determined by minimising the Bayesian Information Criterion (BIC). A power spectral density (PSD) estimate was computed from the resulting AR transfer function on a 4096-point frequency grid, and the frequency of peak power within the high-frequency band (0.15-0.40 Hz) was identified as the participant-specific HF-HRV center frequency.

A participant- and condition-specific bandpass filter was then constructed centered on this peak frequency (± 0.05 Hz), implemented as a zero-phase FIR filter with a Hamming window and filter order set to capture three full cycles of the lower edge. This filter was applied to the original sample-level IBI series (prior to resampling) using zero-phase forward–backward filtering. The instantaneous phase of the filtered signal was then extracted via the Hilbert transform, yielding a continuous HF-HRV phase time series in radians.

### CFA-Targeted ICA

In preparation for a second ICA targeting the cardiac field artifact (CFA) and further downstream analyses, the EEG, ECG, and HF-HRV phase time series were epoched and aligned using the shared light-pulse trigger signal recorded simultaneously on both amplifiers. Epoch boundaries were defined as the interval between successive trigger onsets, anchoring temporal alignment to a common reference regardless of any differences between systems. For each epoch, segments from each data stream were indexed using the synchronized trigger timestamps and trimmed to the shortest of the three to ensure sample-exact alignment before concatenation.

Following synchronization, a second round of ICA was performed to attenuate the CFA following methods described in Corcoran et al. (2025). All data were downsampled to 200 Hz. R-waves were detected from the ECG using the Pan-Tompkins algorithm, with the same two-stage amplitude and physiological bounds guardrail procedure applied as described above. R-wave events were inserted as markers, after which the continuous EEG data were decomposed via ICA. To identify cardiac components, both the ICA activations and the ECG signal were low-pass filtered below 25 Hz using a zero-phase fourth-order Butterworth filter and submitted to the Hilbert transform to obtain instantaneous phase estimates. Component–ECG phase locking was then quantified for each component as the mean resultant length of the phase-angle differences between each component activation and the ECG signal across all samples. Components exceeding 1.96 standard deviations of the participant-level phase-locking distribution were flagged as cardiac in origin. To limit over-rejection, removal was restricted to a maximum of one component per recording. The flagged component was then subtracted from the continuous EEG prior to saving (see Appendix A).

### EEG Oscillatory Amplitude Estimation

The cleaned, synchronized data were screened for artifacts using a sliding non-overlapping 1-second window. Any window in which any EEG channel exceeded ± 50 µV was rejected and excluded from further analysis as well as the corresponding HF-HRV phase window. Amplitude envelopes for theta (4-8 Hz) and alpha (8-12 Hz) were then extracted from the artifact-free data at fronto-central electrode Fz. The signal was bandpass filtered into each frequency band using a zero-phase FIR filter with a Hamming window, with filter order set to capture three full cycles of the lower band edge. The instantaneous amplitude envelope of each filtered signal was extracted via the Hilbert transform.

### Phase-Amplitude Coupling

Phase-amplitude coupling (PAC) was estimated by combining the amplitude envelope of the high-frequency signal with the phase time series of the low-frequency signal into a complex-valued composite signal, whose mean vector length indexed coupling strength (Canolty et al., 2006; Fig. 1H). This particular method, the mean vector length (MVL), is a highly sensitive PAC measure suited for long recordings and monophasic phase-amplitude correspondence^6^ (Hülsemann et al., 2019) –the latter of which was indicated by Sargent et al. (2024).

Empirical MVL was computed separately for HF-HRV phase with theta amplitude and HF-HRV phase with alpha amplitude. To standardize coupling strength, empirical MVL values were referenced to participant-specific surrogate null distributions generated from 1,000 circularly shifted permutations of the high-frequency amplitude time series. For each permutation, the amplitude series was shifted by a random offset of 10-90% of the total series length, after which MVL was recomputed. Empirical MVL was then z-standardized relative to the mean and standard deviation of the corresponding null distribution, yielding standardized MVL values, hereafter referred to as PACz, for theta and alpha bands.

### Extracting Respiratory Descriptives

Because HF-HRV is influenced by respiration, respiratory parameters were extracted to evaluate whether condition-related changes in coupling could be attributable to condition-related changes in breathing. Respiratory effort was processed using the BreathMetrics toolbox (Noto et al., 2018) to derive breathing frequency (Hz) and respiratory depth (a.u.). These measures were retained as candidate covariates in condition-related models.

## Results

Linear regression evaluated the effect of GAD-7 responses on standardized HF-HRV phase-amplitude coupling (PACz) for theta and alpha activity at Fz while controlling for log-transformed EEG band power and HF-HRV power at rest. Bonferroni correction was applied to account for theta and alpha band comparisons. To evaluate spatial specificity, the same regression models were fit separately for theta and alpha activity at Pz, a posterior channel.

Condition-related differences in cardiac, respiratory, and EEG measures were characterized using paired-samples t tests. Condition-related differences in HF-HRV–theta PACz at Fz were then estimated using a mixed-effects model that adjusted for mean heart rate, breathing frequency, and respiratory depth because of their physiological relevance to HF-HRV. Lastly, a GAD-7 by condition interaction was evaluated for HF-HRV-theta PACz at Fz.

Sensitivity analyses using a leave-one-out (LOO) approach are reported in the Supplementary Materials (Appendix B). For each statistically significant effect reported below, the relevant model was re-estimated after sequentially omitting each participant.

### Anxiety Severity Is Associated with Lower Resting-State HF-HRV–Theta PAC

We first tested whether resting-state HF-HRV phase-amplitude coupling at the fronto-central electrode Fz varied with GAD-7 scores while controlling for EEG-band and HF-HRV power (Fig. 2A). The model predicting HF-HRV–theta PACz was significant, F(3, 18) = 4.50, p = 0.016, adjusted R^2^ = 0.33. Higher GAD-7 scores were associated with lower HF-HRV–theta PACz, b = -0.134, 95% CI [-0.213, -0.056], t(18) = -3.59, p_bonf_ = 0.004. By contrast, the corresponding model for HF-HRV–alpha phase-amplitude coupling was not significant (F(3,18) = 0.47, p = 0.708), and GAD-7 was not associated with alpha PACz, b = -0.063, 95% CI [-0.203, 0.077], t(18) = -0.95, p_bonf_ = 0.357.

**Figure 2.**
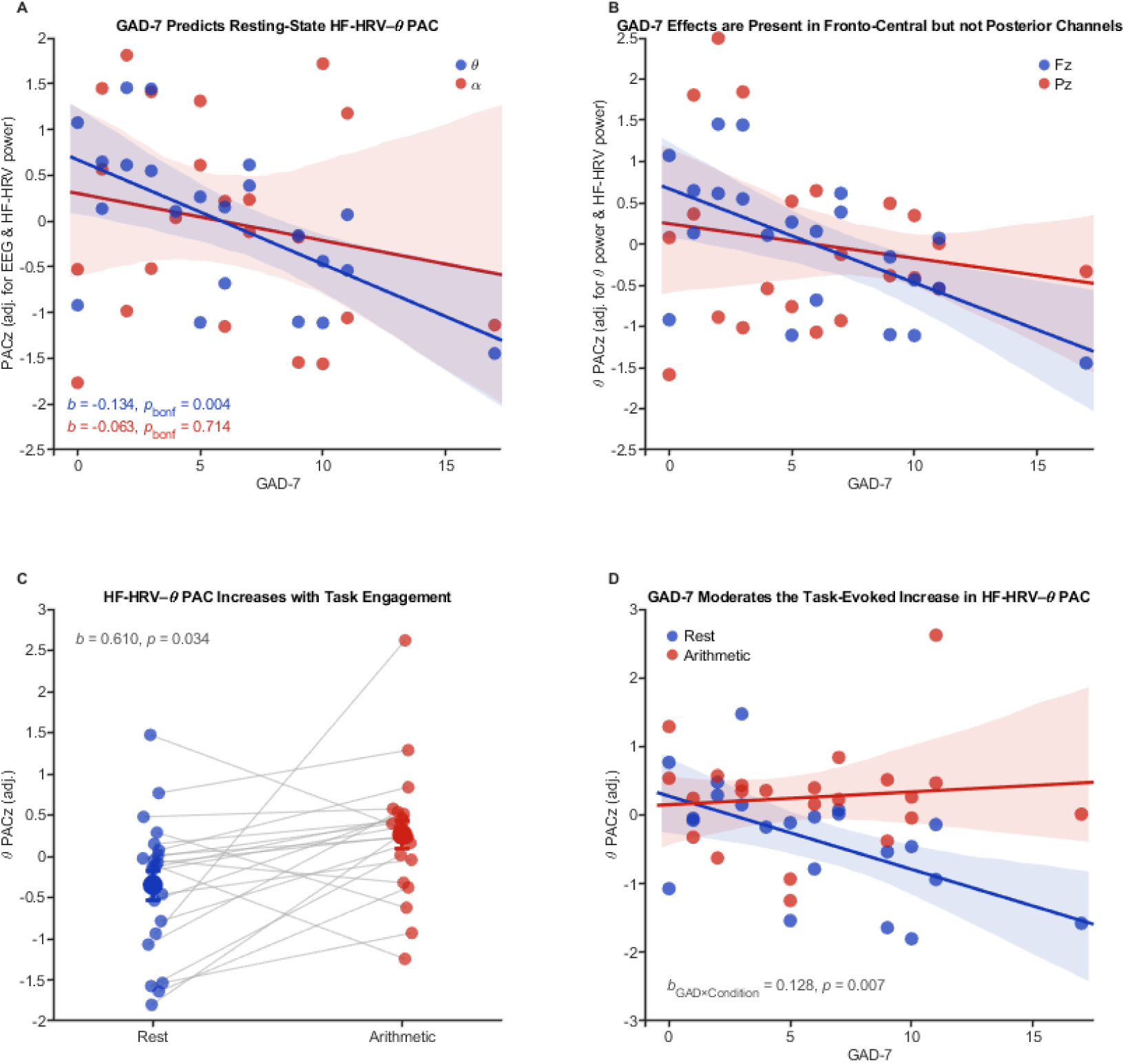
Trait- and State-Related Modulation of HF-HRV–Theta Phase-Amplitude Coupling. *Note. Higher GAD-7 scores were associated with lower resting-state phase-amplitude coupling between HF-HRV phase and theta amplitude at Fz, controlling for log-transformed theta and HF-HRV power (A). The association was observed for the theta band but not the alpha band. The negative association between GAD-7 scores and theta PACz observed at Fz did not generalize to the posterior comparison channel Pz (B). Relative to resting-state, fronto-central HF-HRV–theta PACz increased during a mental arithmetic task while controlling for heart rate, breathing frequency, and respiratory depth (C). GAD-7 scores moderated the condition-related change in HF-HRV–theta PACz: participants with higher anxiety symptomatology showed larger condition-related differences (D)*.

To evaluate whether the theta association generalized to a posterior comparison site, we repeated the model at Pz (Fig. 2B). The posterior theta model was not significant (F(3, 18) = 0.59, p = 0.628), and GAD-7 was unrelated to theta PACz at Pz, b = -0.049, 95% CI [-0.173, 0.074], t(18) = -0.84, p_bonf_ = 0.829. We then formally tested spatial specificity using a mixed-effects model predicting theta PACz from GAD-7, channel (Fz vs. Pz), and log-transformed theta power, including GAD-7 by channel and theta power by channel interactions. The GAD-7 by channel interaction was not significant (b = -0.061, 95% CI [-0.180, 0.058], t(18) = -1.03, p = 0.317). Thus, although the anxiety-related effect was observed at Fz but not Pz, spatial specificity remains descriptive. The posterior alpha model was also not significant (F(3, 18) = 1.32, p = 0.299), and GAD-7 was unrelated to alpha PACz at Pz, b = 0.030, 95% CI [-0.130, 0.191], t(18) = 0.40, p_bonf_ = 1.000.

GAD-7 was not significantly correlated with any conventional autonomic or EEG spectral measure, although higher GAD-7 scores showed marginal negative associations with log-transformed HF-HRV power (r(20) = -0.380, p = 0.081) and RMSSD, r(20) = -0.361, p = 0.099. Given that the hypothesized anxiety-related association was observed only for theta PACz, subsequent condition-related analyses focused on the theta band.

### HF-HRV–Theta PAC Increases During Mental Arithmetic

We first determined what autonomic or neural indices differed between conditions using paired-samples t-tests. Relative to rest, the mental arithmetic task was associated with increases in mean heart rate [M_rest_ = 69.75 bpm, M_arithmetic_ = 72.65 bpm; t(21) = 3.97, p < 0.001] and mean breathing frequency [M_rest_ = 0.18 Hz, M_arithmetic_ = 0.25 Hz; t(21) = 5.63, p < 0.001], as well as decreases in log-transformed LF-HRV power [t(21) = -2.94, p = 0.007] and average respiratory depth [t(21) = -4.17, p < 0.001]. Fz log-transformed theta power [t(21) = -0.856, p = 0.402], HF-HRV power [t(21) = -1.47, p = 0.156], and RMSSD [t(21) = 0.929, p = 0.364] did not differ between conditions.

We next estimated the condition-related differences in fronto-central HF-HRV–theta phase-amplitude coupling while controlling for heart rate, breathing frequency, and respiratory depth (Fig. 2C). The mixed-effects model, including a random intercept, indicated that fronto-central HF-HRV–theta phase-amplitude coupling was significantly higher during arithmetic than rest [b = 0.610, 95% CI [0.048, 1.172], t(39) = 2.20, p = 0.034], independent of cardiac acceleration [b = -0.017, 95% CI [-0.044, 0.010], t(39) = -1.27, p = 0.212], breathing acceleration [b = 2.89, 95% CI [-2.505, 8.285], t(39) = 1.08, p = 0.285], or respiratory depth changes [b = 0.001, 95% CI [0.000, 0.002], t(39) = 1.63, p = 0.111].

### Anxiety Moderates the Task-Evoked Increase in HF-HRV–Theta PAC

We tested whether the task-evoked increase in fronto-central HF-HRV–theta phase-amplitude coupling varied with anxiety symptomatology (Fig. 2D). A mixed-effects model including main effects of GAD-7, heart rate, breathing frequency, respiratory depth, condition, a GAD-7 by condition interaction term, and a random intercept indicated a significant GAD-7 by condition interaction [b = 0.128, 95% CI [0.038, 0.218], t(36) = 2.87, p = 0.007]. Simple-slope analyses indicated that higher GAD-7 scores were associated with lower theta PACz at rest (b = -0.113, 95% CI [-0.188,-0.038], t(37)= -3.06, p= 0.004), but were not associated with theta PACz during mental arithmetic, b = 0.015, 95% CI [-0.058, 0.089], t(37)= 0.42, p= 0.679.

As a sensitivity analysis, we repeated the interaction model while adjusting for log-transformed theta power and HF-HRV power, mirroring the resting-state model. The GAD-7 by condition interaction remained significant, b = 0.129, 95% CI [0.036, 0.223], t(36) = 2.80, p = 0.008.

## Discussion

There is broad and growing interest in linking peripheral bodily measures to mental states because the brain and its cognitions exist within a dynamic somatic environment whose maintenance of homeostasis may exert a strong influence on perception and cognition (Kluger et al., 2024). Heart rate variability (HRV), and particularly high-frequency HRV (HF-HRV), has commanded considerable attention as a peripheral index of autonomic regulation that covaries with mental and physical outcomes. It has been proposed that HF-HRV may covary with regulatory processes because it serves as a scaffold for organizing neural oscillations within prefrontal cortex (Mather & Thayer, 2018). The present study examined whether resting-state phase-amplitude coupling (PAC) between cardiac and neural rhythms varies dimensionally with anxiety symptomatology, and whether heart-brain coupling differs across cognitive states.

Higher GAD-7 scores were associated with reduced coupling between HF-HRV phase and fronto-central theta amplitude while controlling for theta and HF-HRV power. The corresponding analysis at a posterior channel was not significant, suggesting that this association may be more prominent fronto-centrally. Importantly, anxiety symptomatology was unrelated to conventional autonomic and spectral indices, including HF-HRV power, LF-HRV power, LF/HF ratio, RMSSD, mean heart rate, EEG band power, breathing frequency, and respiratory depth.

The anxiety-related decrease in HF-HRV–theta PAC mirrors the reduction reported in individuals with schizophrenia (Sargent et al., 2025), but offers additional nuance. Given that anxiety has been associated with enhanced interoceptive sensitivity (Young et al., 2026), whereas schizophrenia is associated with interoceptive deficits (Koreki et al., 2021; Fan et al., 2025), a simple account in which greater HF-HRV–theta coupling indexes greater interoceptive processing would predict increased, rather than decreased, coupling with anxiety symptomatology. The present results are less consistent with that account. Instead, the similar association with anxiety severity observed here and the reduction reported in individuals with schizophrenia suggest that HF-HRV–theta PAC may reflect a non-interoceptive faculty.

Consistent with this interpretation, fronto-central HF-HRV–theta PAC increased during a (non-interoceptive) cognitive task. Moreover, anxiety symptomatology moderated this task-related increase. Simple-slope analyses indicated that greater anxiety was associated with lower coupling at rest but not during mental arithmetic, suggesting that task engagement attenuated the anxiety-related reduction in resting-state HRV–EEG coupling.

### An Inhibition-Related Account

One candidate, though speculative, interpretation of our findings is that fronto-central HF-HRV–theta PAC indexes a state-sensitive component of inhibitory control. The neurovisceral integration model provides a foundation for this account by linking cardiac variability to the central autonomic network (CAN; Thayer & Lane, 2000). Efferent projections from the CAN innervate the heart, while several of its cortical nodes—including orbitofrontal, ventromedial prefrontal, and anterior cingulate cortices—are implicated in cognitive inhibition (Thayer et al., 2009), a common factor underlying executive function (Miyake & Friedman, 2012). Consistent with this framework, a recent meta-analysis found that HF-HRV was associated with all three executive function factors, but more strongly with cognitive inhibition and cognitive flexibility than with working memory updating (Magnon et al., 2022). It is likewise notable that the present association was observed for theta but not alpha, as theta activity is characteristically prominent over fronto-central sites (Donoghue et al., 2020) and fronto-central theta has been implicated in signaling the need for and coordinating the implementation of executive functions (Cavanagh & Frank, 2014).

Anxiety has been associated with impaired inhibitory control across behavioral and neural measures (Xia et al., 2020; Wang et al., 2024), but anxious individuals derive disproportionate benefit from inhibitory control interventions (Mamat & Anderson, 2023). If HF-HRV–theta PAC indexes inhibitory control, the present findings align with this pattern: greater anxiety was associated with lower coupling at rest, but this difference was attenuated during mental arithmetic, a task that places demands on executive control. This account remains conjectural because we did not directly measure inhibitory control, but it generates actionable hypotheses.

Future work would benefit from incorporating a direct measure of inhibitory control, such as the think/no-think paradigm (Anderson & Green, 2001), to evaluate the proposed inhibition-related interpretation of fronto-central HF-HRV–theta PAC. Incorporating an interoceptive task, such as the Heartbeat Counting (Schandry, 1981) or Heartbeat Detection tasks (Ring & Brener, 2018), would clarify whether HF-HRV–theta PAC is related to interoceptive processing, a non-interoceptive regulatory process, or both.

## Limitations

Several limitations constrain interpretation of the present findings. First, the small, nonclinical sample limits inferences regarding Generalized Anxiety Disorder. Second, the association between anxiety symptomatology and HF-HRV–theta PAC was cross-sectional, precluding conclusions regarding causal direction. Anxious states may attenuate resting-state coupling or attenuated resting-state coupling may increase one’s vulnerability to anxious states. Lastly, resting-state and mental arithmetic conditions differed in duration. We mitigated this potential confound by standardizing each empirical PAC estimate against a condition-specific surrogate distribution generated from a time series of equivalent length.

## Conclusion

In conclusion, anxiety symptomatology was associated with fronto-central HF-HRV–theta phase-amplitude coupling in a manner not captured by conventional autonomic or neural measures alone. Greater anxiety was associated with reduced coupling at rest, while mental arithmetic increased coupling, with larger increases observed among more anxious individuals. The present association with anxiety severity parallels the reduction in fronto-central HF-HRV–theta coupling reported in schizophrenia, despite evidence that anxiety and schizophrenia are associated with opposing patterns of interoceptive sensitivity (Fan et al., 2025; Young et al., 2026). Together, these findings suggest that this coupling may not be monotonically related to interoceptive processing. Instead, HF-HRV–theta PAC may covary with a state-sensitive regulatory process whose expression differs between unconstrained rest and constrained task engagement.

## Research Transparency Statement

### General Disclosure

All authors declare they have no conflicts of interest.

### Funding

As secondary analyses on an existing dataset, the authors report no funding involved in the present study.

### Open Practices

No aspects of the study were preregistered. All analytic code and (raw and processed) anonymized data can be made available upon request to the first author.

### Ethics

The experiment was approved by the University of California, Santa Barbara’s Institutional Review Board under protocol number 202-23-0348.

## Supplementary Materials

### Appendix A. CFA-Targeted ICA in a Representative Participant

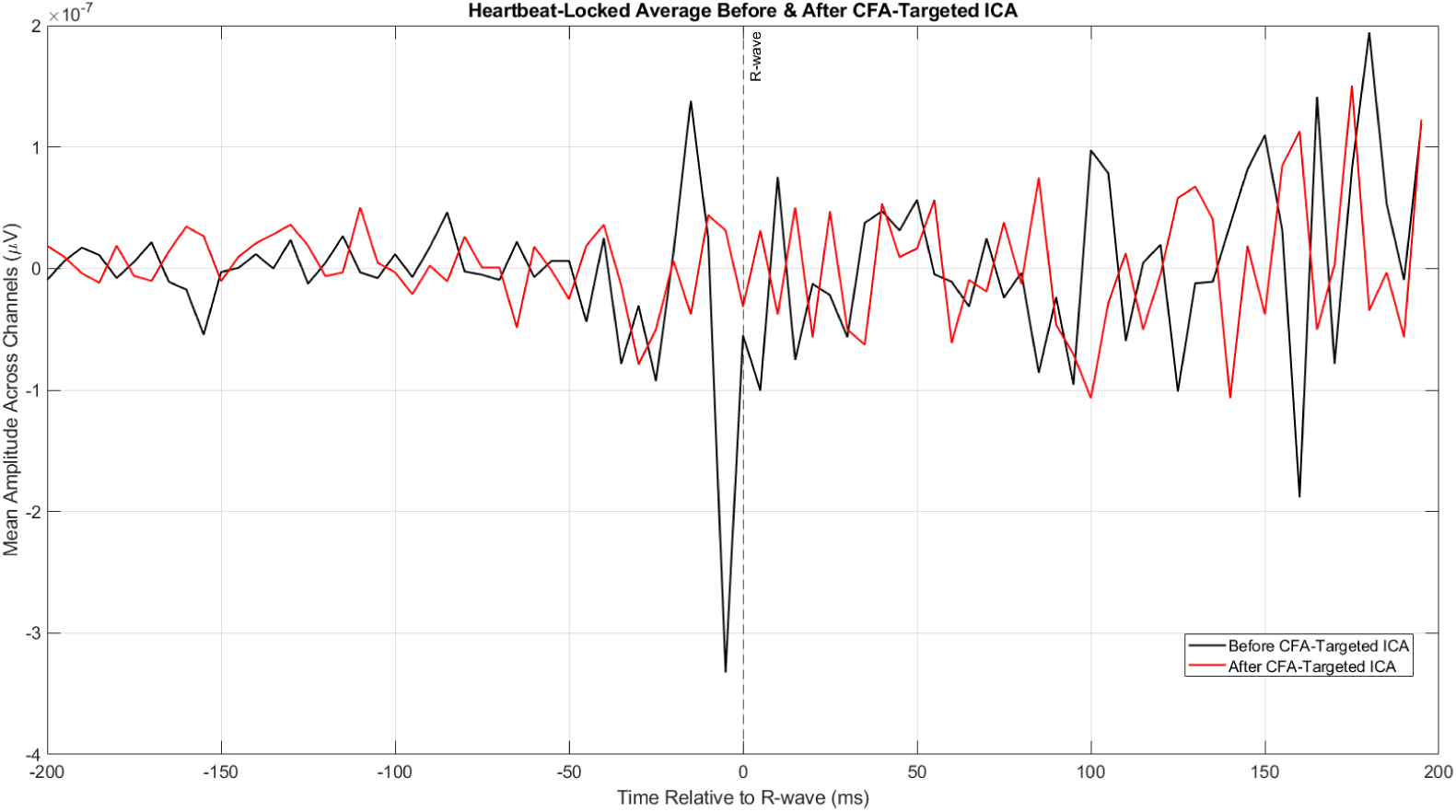
*Note. To attenuate cardiac contamination of the EEG, including the cardiac field artifact (CFA) arising from volume-conducted cardiac electrical activity, we conducted a second, CFA-targeted independent component analysis (ICA). This figure illustrates the effect of this procedure in a representative participant. Before ICA-based removal, the heartbeat-locked EEG average showed a prominent deflection contemporaneous with the ECG R-wave, consistent with CFA. After removal of the component flagged as cardiac in origin, this R-wave–locked deflection was substantially attenuated. Importantly, this peri-R-wave deflection should not be interpreted as a heartbeat-evoked potential (HEP). Although HEPs are also time-locked to the R-wave, their putatively interoceptive components are examined in later post-R-wave intervals, approximately 150–500 ms after the heartbeat*.

### Appendix B. Leave-One-Out Sensitivity Analysis

To assess whether statistically significant theta-band findings were disproportionately influenced by any individual participant, we conducted leave-one-out (LOO) sensitivity analyses for each primary model. Each model was re-estimated 22 times, sequentially omitting one participant on each iteration. We examined the stability of the coefficient direction, coefficient magnitude, and statistical significance of the focal effect.

The resting-state GAD-7 effect and the GAD-7 by condition interaction remained statistically significant across all 22 iterations. The main effect of condition remained positive across all iterations but was statistically significant in 15 of 22 models, indicating greater sensitivity to individual observations. For the resting-state Fz regression predicting HF-HRV–theta PACz from GAD-7 while controlling for log-transformed theta and HF-HRV power, the GAD-7 coefficient remained negative, ranging from -0.151 to -0.121, with p values ranging from < 0.001 to 0.013. For the mixed-effects model testing the condition effect, the condition coefficient remained positive, ranging from 0.289 to 0.741, with p values ranging from 0.012 to 0.222. Finally, for the mixed-effects model testing the GAD-7 by condition interaction, the interaction coefficient remained positive, ranging from 0.102 to 0.166, with p values ranging from < 0.001 to 0.016.

## Footnotes

1 Averaged Fz, FCz, F1, F2, and AFz.

2 For example, mutual information or transfer entropy (see Candia-Rivera et al., 2025).

3 The two excluded participants had anomalous resting-state RMSSD values of 380.543 and 285.463 ms. Among the remaining participants, resting-state RMSSD ranged from a typical 10.200 to 84.119 ms.

4 Gain: 1000. Mode: Norm. 35 Hz LPN: On. HP: 0.05 Hz.

5 Gain: 10. LP: 1 Hz. HP: DC.

6 Monophasic PAC describes a correspondence in which amplitude is preferentially aligned to one portion of phase (e.g., peak amplitude at phasic troughs) while biphasic PAC describes a correspondence in which amplitude is, for example, preferentially aligned to both phasic peaks and troughs.

## Notes

### Competing Interest Statement

The authors have declared no competing interest.

