## Supplementary Materials for "Resting-state and task-evoked phase-amplitude coupling between cardiac and neural rhythms is sensitive to anxiety severity"

**Appendix A.** CFA-Targeted ICA in a Representative Participant.

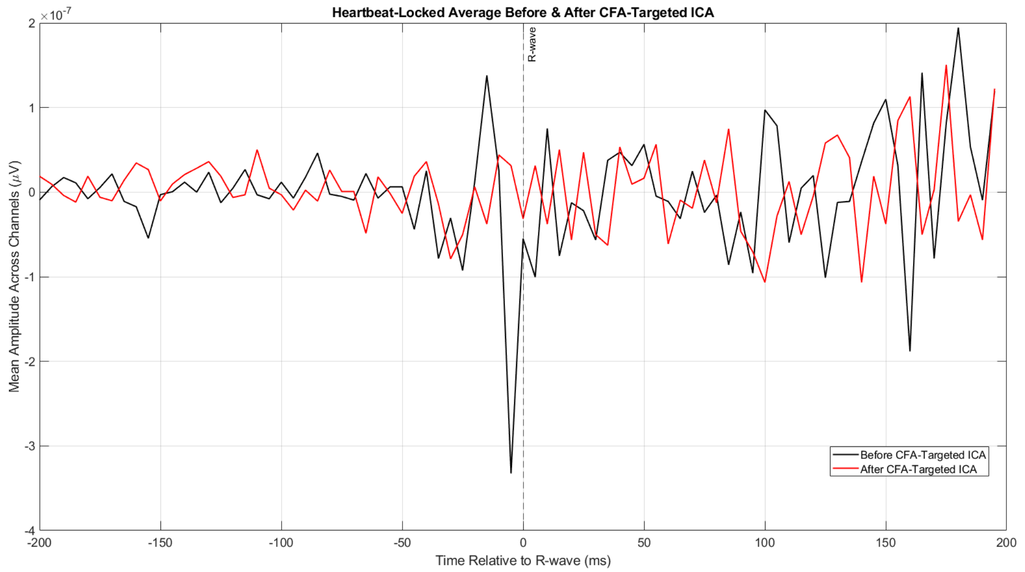


*Note. To attenuate cardiac contamination of the EEG, including the cardiac field artifact (CFA) arising from volume-conducted cardiac electrical activity, we conducted a second, CFA-targeted independent component analysis (ICA). This figure illustrates the effect of this procedure in a representative participant. Before ICA-based removal, the heartbeat-locked EEG average showed a prominent deflection contemporaneous with the ECG R-wave, consistent with CFA. After removal of the component flagged as cardiac in origin, this R-wave–locked deflection was substantially attenuated. Importantly, this peri-R-wave deflection should not be interpreted as a heartbeat-evoked potential (HEP). Although HEPs are also time-locked to the R-wave, their putatively interoceptive components are examined in later post-R-wave intervals, approximately 150–500 ms after the heartbeat.*

**Appendix B.** Leave-One-Out Sensitivity Analysis.

To assess whether statistically significant theta-band findings were disproportionately influenced by any individual participant, we conducted leave-one-out (LOO) sensitivity analyses for each primary model. Each model was re-estimated 22 times, sequentially omitting one participant on each iteration. We examined the stability of the coefficient direction, coefficient magnitude, and statistical significance of the focal effect.

The resting-state GAD-7 effect and the GAD-7 by condition interaction remained statistically significant across all 22 iterations. The main effect of condition remained positive across all iterations but was statistically significant in 15 of 22 models, indicating greater sensitivity to individual observations. For the resting-state Fz regression predicting HF-HRV–theta PACz from GAD-7 while controlling for log-transformed theta and HF-HRV power, the GAD-7 coefficient remained negative, ranging from -0.151 to -0.121, with p values ranging from < 0.001 to 0.013. For the mixed-effects model testing the condition effect, the condition coefficient remained positive, ranging from 0.289 to 0.741, with p values ranging from 0.012 to 0.222. Finally, for the mixed-effects model testing the GAD-7 by condition interaction, the interaction coefficient remained positive, ranging from 0.102 to 0.166, with p values ranging from < 0.001 to 0.016.
